# Optimizing DNA extraction methods for Nanopore sequencing of *Neisseria gonorrhoeae* direct from urine samples

**DOI:** 10.1101/827485

**Authors:** Teresa L. Street, Leanne Barker, Nicholas D. Sanderson, James Kavanagh, Sarah Hoosdally, Kevin Cole, Robert Newnham, Mathyruban Selvaratnam, Monique Andersson, Martin J. Llewelyn, Justin O’Grady, Derrick W. Crook, David W. Eyre, the GonFast Investigators Group, Joanna Rees, Emily Lord, Suneeta Soni, Celia Richardson, Joanne Jessop, Tanya Adams

## Abstract

**Background:** Empirical gonorrhoea treatment at initial diagnosis reduces onward transmission. However, increasing resistance to multiple antibiotics may necessitate waiting for culture-based diagnostics to select an effective treatment. There is a need for same-day culture-free diagnostics that identify infection and detect antimicrobial resistance.

**Methods:** We investigated if Nanopore sequencing can detect sufficient *N. gonorrhoeae* DNA to reconstruct whole genomes directly from urine samples. We used *N. gonorrhoeae* spiked urine samples and samples from gonorrhoea infections to determine optimal DNA extraction methods that maximize the amount of *N. gonorrhoeae* DNA sequenced whilst minimizing contaminating host DNA.

**Results:** In simulated infections the Qiagen UCP Pathogen Mini kit provided the highest ratio *N. gonorrhoeae* to human DNA and the most consistent results. Depletion of human DNA with saponin increased *N. gonorrhoeae* yields in simulated infections, but decreased yields in clinical samples. In ten urine samples from men with symptomatic urethral gonorrhoea, ≥87% coverage of an *N. gonorrhoeae* reference genome was achieved in all samples, with ≥92% coverage breath at ≥10-fold depth in 7 (70%) samples. In simulated infections if ≥10^4^ CFU/ml of *N. gonorrhoeae* was present, sequencing of the large majority of the genome was frequently achieved. *N. gonorrhoeae* could also be detected from urine in cobas PCR Media tubes and from urethral swabs, and in the presence of simulated *Chlamydia* co-infection.

**Conclusion:** Using Nanopore sequencing of urine samples from men with urethral gonorrhoea sufficient data can be obtained to reconstruct whole genomes in the majority of samples without the need for culture.

## Introduction

Multidrug resistant *N. gonorrhoeae* infection is a substantial public health threat.^1,2^ To reduce the spread of antimicrobial resistance, empirical dual therapy with single-dose azithromycin and ceftriaxone, the last two mainstream treatment options, is widely recommended.^2^ However, increasing azithromycin resistance potentially undermines this approach leaving ceftriaxone empirical monotherapy as a last option,^3^ but with ceftriaxone resistant cases recently reported in several countries worldwide.^4,5^

Single dose gonorrhoea treatment at the point of initial diagnosis reduces onward transmission.^6^ However, rising resistance rates can necessitate delays of up to several days while culture-based susceptibility testing is performed to direct effective treatment, potentially increasing transmission of resistant organisms. Therefore, there is a need for same-day culture-free diagnostics that are able to confirm both the presence of infection and detect antimicrobial resistance.^7^

Clinical metagenomics has the potential to detect *N. gonorrhoeae* and antimicrobial resistance determinants via direct sequencing of all DNA present in a clinical sample.^8^ Previous studies have shown this is possible for *N. gonorrhoeae* using Illumina sequencing,^9^ but this approach is not suitable for rapid near-patient deployment. Furthermore, current approaches are limited by obtaining sufficient pathogen DNA against a background of human host DNA and DNA from other bacteria.^10^ A range of approaches have been used to increase pathogen DNA yields, e.g. microbial enrichment via immunomagnetic separation^11^ for sequencing *Chlamydia trachomatis* from urine, or capture of pathogen DNA after DNA extraction using RNA baits.^12^ Several commercial kits exist for selective depletion of human DNA.^10^ Despite this, obtaining sufficient pathogen DNA for whole-genome reconstruction from metagenomic sequencing remains challenging and the short reads generated by Illumina sequencing makes it difficult to accurately assign resistance determinants to specific species in metagenomic samples.

The Oxford Nanopore sequencing platform has the potential to overcome these challenges and deliver *N. gonorrhoeae* and antimicrobial resistance detection in a benchtop format that yields results within a few hours. The long-reads generated have the potential to link antimicrobial resistance determinants accurately to given species as greater genetic context is provided. Here we build on previous work applying Nanopore sequencing for diagnosis of urinary tract infection^13^ and develop optimized laboratory protocols for sequencing *N. gonorrhoeae* directly from patient urine samples.

## Materials and Methods

### Samples

Urine samples collected as part of routine clinical testing of patients at sexual health clinics in Oxfordshire, UK were obtained from the Microbiology Laboratory of Oxford University Hospitals and used for initial method development. Samples were tested for *N. gonorrhoeae* and *C. trachomatis* using the BD Viper system with confirmatory testing for *N. gonorrhoeae* undertaken using the BD Max platform (Becton Dickinson, Wokingham, Berkshire, UK). Samples, that would otherwise have been discarded, were obtained for research use after completion of routine testing.

Additionally, samples were collected from participants recruited at Sexual Health Clinics at the Churchill Hospital, Oxford University Hospitals NHS Foundation Trust, and Brighton and Sussex University Hospitals NHS Trust. Male patients presenting with symptomatic urethritis were eligible to participate and were recruited following informed consent. In this current study, urine samples from participants recruited in Brighton were used to test the performance of metagenomic sequencing in clinical infection. Urine samples were collected into universal tubes containing boric acid (Medline Scientific, Chalgrove, Oxfordshire, UK), and latterly also into cobas PCR Media tubes (Roche Molecular Systems, Pleasanton, CA, USA), for stabilization during transportation to Oxford. Urethral swabs were collected and placed into Sigma VCM preservation medium (MWE, Corsham, Wiltshire, UK). This study was conducted with NHS Research Ethics approval (reference 19/EM/0029).

### Human DNA depletion and microbial DNA extraction

To optimize laboratory methods for the selective extraction of high-quality, long-fragment *N. gonorrhoeae* DNA directly from urine we tested 16 approaches: four DNA extraction methods in combination with either no human cell/DNA depletion or one of three human cell/DNA depletion protocols (Figure S1). For each approach, *N. gonorrhoeae* nucleic acid amplification test (NAAT)-negative urines were spiked at 10^3^, 10^5^, 10^7^ colony forming units (CFU)/ml with a *N. gonorrhoeae* reference strain. Testing at each dilution was undertaken in triplicate using three *N. gonorrhoeae* reference strains, WHO F, WHO V, and WHO X (obtained from Public Health England’s National Collection of Type Cultures), and a starting urine volume of 3ml. Multiple negative urine samples were pooled to allow the same baseline urine to be tested across each depletion and extraction protocol. The number of CFU/ml present in each spike was initially determined using dilutions of a 0.5 McFarland standard; the resulting dilutions were cultured at 37°C with 5% CO_2_ for 24 hours on Lysed GC Selective Agar (Oxoid, Basingstoke, Hampshire, UK), to enable the final *N. gonorrhoeae* CFU/ml achieved to be measured.

Three human cell/DNA depletion methods were tested: differential centrifugation; saponin-based differential lysis followed by nuclease digestion; and the MolYsis Basic5 kit (Molzym, Bremen, Germany). Urine samples were also processed with no human DNA depletion, as a negative control. Differential centrifugation was performed as previously described.^13^ The bacterial cell pellet was washed with 1 ml PBS before proceeding to DNA extraction. Saponin-based differential lysis and nuclease digestion was performed as previously described.^14^ Human DNA depletion and enrichment of microbial DNA with the MolYsis Basic5 kit was performed per the manufacturer’s instructions.

The four DNA extraction methods assessed included mechanical lysis followed by ethanol precipitation, as described previously;^15^ and the MagMAX Total Nucleic Acid Isolation kit (ThermoFisher Scientific, Waltham, MA, USA); QIAamp UCP Pathogen Mini kit (Qiagen, Hilden, Germany); and i-genomic Urine DNA Extraction Mini kit (iNtRON Bio, Burlington, MA, USA), all performed following the manufacturer’s instructions. Extracted DNA was purified using AMPure XP solid-phase reversible immobilization (SPRI) beads (Beckman Coulter, High Wycombe, UK), eluted in 26 µl of TE buffer and quantified using a Qubit 2.0 fluorometer (Life Technologies, Paisley, UK).

### PCR analysis of *N. gonorrhoeae* and human DNA

Quantitative real-time PCR (qPCR) was performed to determine the relative amounts of both *N. gonorrhoeae* and human DNA in the DNA extracts from the initial laboratory optimization methods. qPCR was performed on a Stratagene MX3005P QPCR System (Agilent Technologies, Santa Clara, CA, USA) using Luna Universal Probe qPCR Master Mix (New England Biolabs, Ipswich, MA, USA). Primers and probes were used to target the β-actin gene for human DNA detection^16^ and the *porA* pseudogene for detection of *N. gonorrhoeae* (papTM).^17^ Reactions were performed in 20 μl with 2 µl of template DNA, 0.4 μM of each primer and 0.2 μM of the probe. Cycling conditions were an initial denaturation at 95 °C for 1 min, followed by 40 cycles of 95 °C denaturation for 15 s and 60 °C extension for 30 s. *N. gonorrhoeae* genomic DNA, extracted from cultures of WHO F, WHO V and WHO X reference strains, was diluted to 100,000 genome copies per μl then serially diluted to 10 genome copies per μl and used to create copy number standard curves. Human genomic DNA (Promega, Madison, WI, USA) was diluted to 10,000 genome copies per μl then serially diluted to 10 genome copies per μl and used to create a human DNA copy number standard curve. Negative controls, replacing template DNA with water, were also performed. All qPCR assays were performed in triplicate and the mean value used in analyses.

### In-depth spiking and metagenomic whole genome sequencing

Using the optimal laboratory method determined in the spiking experiments, an extended set of spiking experiments were performed to determine the limit of detection of this protocol for *N. gonorrhoeae* DNA. 3 ml of *N. gonorrhoeae* NAAT-negative pooled urine, was spiked with one of the WHO F, WHO V or WHO X *N. gonorrhoeae* reference strains, using dilutions of a 0.5 McFarland standard, to target 10^1^, 10^2^, 10^3^, 10^4^, 10^5^ or 10^6^ CFU/ml, using the three reference strains as replicates (n=19 including an un-spiked urine sample as a negative control). DNA extracts were assessed for human and microbial DNA content by metagenomic sequencing. Libraries were prepared for sequencing on an Oxford Nanopore GridION (Oxford Nanopore Technologies (ONT), Oxford, UK) using the Rapid PCR Barcoding kit (SQK-RPB004) (ONT), with modifications to the manufacturers’ protocol as described by Charalampous *et al*.^14^ Briefly, up to 10 ng of input DNA and 2.5 μl of FRM (Fragmentation Mix) were used in the tagmentation reaction with a final volume of 10 μl. For samples that were not able to achieve 10 ng, a total volume of 7.5 μl was used in the tagmentation reaction regardless of the quantity of DNA this represented. The PCR reaction was performed in a double volume of 100 μl, with 25 cycles and an elongation time of 6 minutes. Post-PCR, DNA was purified with AMPure XP SPRI beads (Beckman Coulter, High Wycombe, UK) and eluted in 10 μl. Initially, all samples were sequenced individually on ONT FLO-MIN106D (v.R9.4.1) flowcells regardless of the quantity of DNA after PCR. Subsequently, only samples with amounts of DNA quantifiable by a Qubit fluorometer after PCR were carried forwards for sequencing. Between 3 and 85 fmol of library were loaded per flowcell. One sample was run per flow cell for all experiments.

### DNA extraction in simulated co-infection

To assess sequencing in the presence of co-infection, Chlamydia NAAT-positive urine collected from three patients was used. 10^2^ and 10^4^ CFU/ml of *N. gonorrhoeae* WHO F reference strain was individually spiked into 3 ml of urine from each patient. This was repeated for *N. gonorrhoeae* WHO X and WHO V and an un-spiked sample from each patient was used as a negative control (n=18). Following spiking, each sample was split and DNA was extracted with either the optimal extraction protocol of saponin-based differential lysis and the QIAamp kit, or with saponin-based differential lysis followed by mechanical lysis and ethanol precipitation (to ascertain whether it was possible to recover *N. gonorrhoeae* and *C. trachomatis* DNA, and to assess any potential bias in DNA recovery between the two methods). Human and microbial DNA content were assessed by sequencing on an ONT GridION, as described above.

### DNA extraction from NAAT-positive urine samples and urethral swabs

Samples from study participants were used to assess the real-world performance of our methods in *N. gonorrhoeae* NAAT-positive urine samples from men with urethral gonorrhoea infection. Clinical samples were tested with the FTD Gonorrhoea confirmation NAAT assay (Fast Track Diagnostics, Sliema, Malta). DNA was extracted from 3 ml of urine from ten individual participants using the QIAamp UCP Pathogen Mini kit (chosen on the basis of the simulated infection experiments) with and without saponin-based human DNA depletion. DNA was sequenced on an ONT GridION as described above, with the DNA extracted from urine both with and without saponin multiplexed as a pair on a single flowcell. Urine from four of the 10 individual participants was also collected into cobas PCR Media tubes designed for diagnostic molecular *N. gonorrhoeae* and *C. trachomatis* testing. DNA was extracted from 4 ml obtained from these tubes using the QIAamp kit (without saponin treatment) and with or without a prior mechanical lysis step using bead beating as previously described,^15^ with the resulting DNA extracts from a single sample multiplexed as a pair on a single flowcell. Urethral swabs were obtained from nine of the participants and stored in Sigma VCM preservation medium for transport. DNA was extracted from 3ml of this medium, following vortexing for 30 seconds, using the QIAamp UCP Pathogen Mini kit without saponin treatment and sequenced as described above with a single sample per flowcell.

### Bioinformatics analysis

Nanopore sequences were basecalled using Guppy version 1.8.5 (ONT) and demultiplexed with Porechop 0.2.4 (https://github.com/rrwick/Porechop) using the default settings. Reads were taxonomically classified using Centrifuge^18^ using a database of NCBI RefSeq bacterial and viral genomes submitted by August 10, 2018 and the human reference genome (GRCh38). Reads classified as human were securely deleted prior to subsequent analyses. During the study, to reduce the number of reads not assigned a barcode following demultiplexing, sequenced data from samples obtained from participants were basecalled and demultiplexed again using an updated version of Guppy (3.3.0+ef22818, ONT). Basecalling was done with the HAC (“high accuracy”) model and recommended kit and flowcell configurations. Barcode demultiplexing was done with default parameters. As all human DNA sequenced was removed prior to repeat basecalling/demultiplexing, analyses of the relative proportion of human to bacterial DNA sequenced are based on the data demultiplexed with Porechop.

Classified reads were aligned to reference genomes using Minimap2^19^ and filtered to a mapQ of 50 by samtools^20^ as described in the CRuMPIT workflow^21^ (https://gitlab.com/ModernisingMedicalMicrobiology/CRuMPIT). CRuMPIT was used to generate per species metrics for read numbers, bases, and coverage depth and breadth. Within CRuMPIT the depth and breadth of the mapped reads were assessed using samtools.^20^ Additional plots were generated using the following python libraries: pandas, seaborn and matplotlib. The proportion of the *Neisseria gonorrhoeae* NCCP11945 (Accession NC_011035.1) and *Chlamydia trachomatis* D/UW-3/CX (Accession NC_000117.1) reference genomes covered at ≥10-fold coverage is reported as a summary of the proportion of the genome likely to have informative coverage given the inherent ~10% base inaccuracy of ONT reads.

### Data availability

Nanopore sequence read data generated from simulated and clinical infections are available from NCBI/EBI under study accession number PRJEB35173.

## Results

### *N. gonorrhoeae* DNA extraction and human DNA depletion

The 4 DNA extraction methods tested in simulated infections yielded varying amounts of *N. gonorrhoeae* DNA. For example, in *N. gonorrhoeae* NAAT-negative urines spiked with 10^5^ CFU/ml of an *N. gonorrhoeae* reference strain, ethanol precipitation and the MagMAX kit yielded higher amounts of *N. gonorrhoeae* DNA: samples contained a median (interquartile range, IQR) of 695 (60-2020), 318 (68-595), 588 (16-1037), 180 (41-241) copies/ml of *porA* across all human DNA depletion methods using ethanol precipitation, i-genomic, MagMAX and QIAamp kits respectively. Figure S2A shows *porA* qPCR for all spike concentrations tested. The amount of hands on time and total time taken for each of the four DNA extraction methods and their approximate cost is shown in Table S1.

Across all DNA extraction methods, the MolYsis Basic5 kit or saponin-based differential lysis successfully depleted the most human DNA when compared to the control samples without human DNA depletion. The median (IQR) percentage of human DNA present before treatment remaining after depletion was 0.3% (<0.1-0.7%) and 1.1% (0.4-2.3%) respectively. Differential centrifugation did not lead to any observable depletion of human DNA, and instead enriched for human DNA, 154% (108-277%) (Figure S2B).

Saponin-based differential lysis produced the highest ratio of *N. gonorrhoeae* to human DNA across all the spikes and for all the DNA extraction protocols tested, with little observable difference between the four extraction methods (Figure S2C). The QIAamp kit produced a higher ratio of *N. gonorrhoeae* to human DNA at the lowest spiked amount and the most consistent results overall. From these results and considering ease of sample processing within the laboratory with each of the protocols tested, we chose to use the saponin-based differential lysis and nuclease digestion followed by the QIAamp UCP Pathogen Mini Kit as our laboratory method for subsequent experiments.

### Limit of detection

For these experiments, the concentration of *N. gonorrhoeae* reference strain spikes achieved ranged from approximately 10^1^ to 10^6^ CFU/ml (see supplementary Figure S3 for comparison of target and actual spiking concentrations). Samples generated between 4 and 13 gigabases of taxonomically classified sequence data (with the exception of one sample that failed library preparation and generated no sequence data). The majority of reads were classified as bacterial in all samples sequenced (Figure 1A, see supplementary Figure S4 for a breakdown of the most common bacterial species identified). There was no relationship between the total number of bacterial reads and the concentration of the *N. gonorrhoeae* spike. This reflects the high concentrations of background bacteria present, such that at low spike concentrations the bacteria sequenced were predominantly other species, whereas at higher spikes *N. gonorrhoeae* sequence dominated (Figure 1B). The proportion of reads classified as human was <5% for all samples, and <1.5% for 10/18 samples. Bases classified as *N. gonorrhoeae* were detected in all samples spiked at ≥10^3^ CFU/ml (Figure 1B). On mapping, we observed coverage at ≥10-fold to ≥75% of the NCCP11945 reference genome in all samples spiked with ≥10^4^ CFU/ml (Figure 1C). Samples spiked at ≥10^5^ CFU/ml achieved a median (IQR) coverage breadth of 98% (97.2-98.1) of the *N. gonorrhoeae* reference genome at an average coverage depth of 2730 (1133-4027). We observed reads classified as *N. gonorrhoeae* in our un-spiked negative control urine sample, which mapped to <14% of the whole reference genome at a depth of ≥10-fold (Figure S5). However, no similar contamination was seen in six subsequent negative urine sample sequences (Figure S6B).

**Figure 1.**
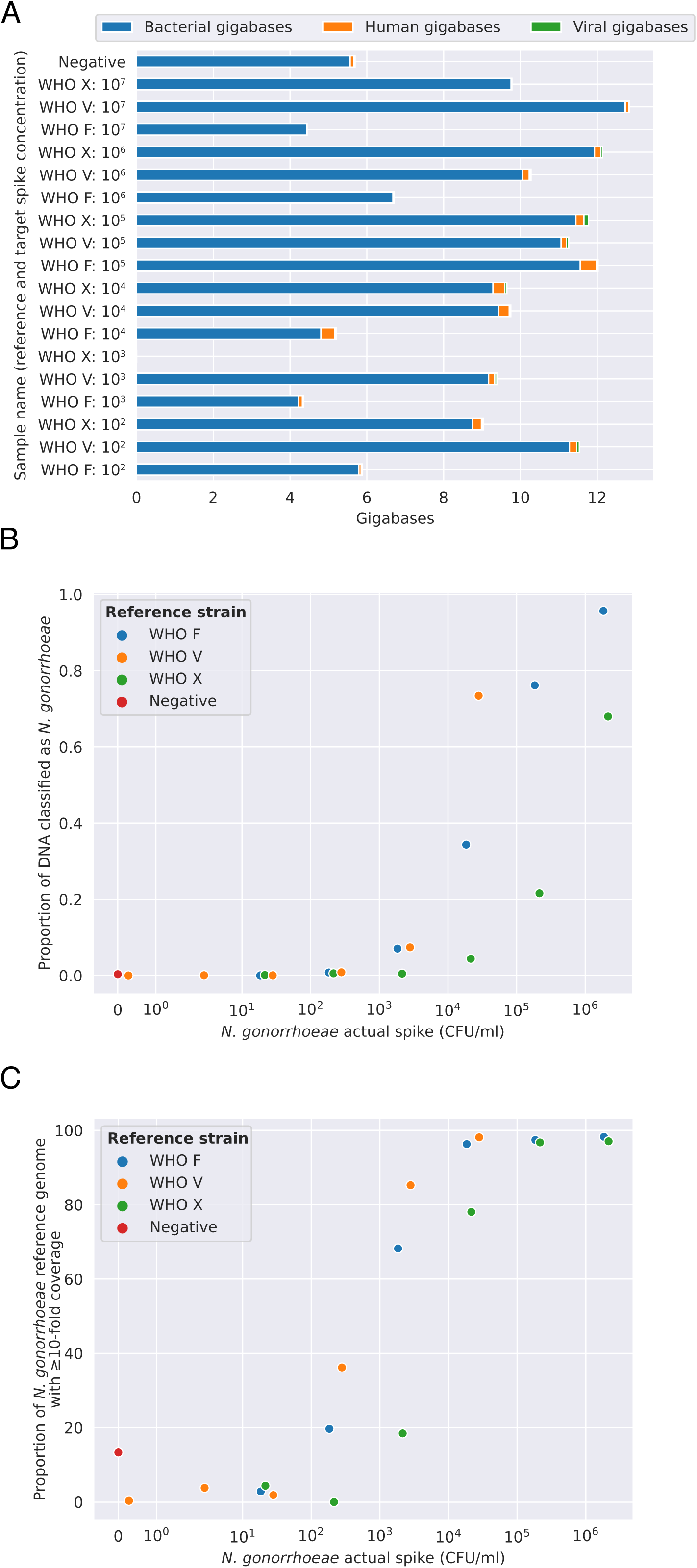
*N. gonorrhoeae* simulated infections: limit of detection using Nanopore sequencing. Panel A shows the proportion of sequenced reads classified as human, bacterial or viral. Panel B shows the proportion of bacterial bases classified as *N. gonorrhoeae*, and panel C the proportion of the NCCP11945 *N. gonorrhoeae* reference genome covered at ≥10-fold depth. For WHO V (orange markers) the actual spike concentration achieved was lower than the target (see Figure S3).

### Detection of *N. gonorrhoeae* in the presence of *Chlamydia trachomatis*

Our protocol also allowed direct detection of both *N. gonorrhoeae* and *C. trachomatis* DNA in simulated co-infections. DNA from Chlamydia NAAT-positive urine spiked with *N. gonorrhoeae* was extracted by using the QIAamp kit or mechanical lysis followed by ethanol precipitation, both following saponin treatment. Approximately 1.2 to 12.5 gigabases of taxonomically classified sequence data were generated from 16/18 samples sequenced: two samples performed sub-optimally in the library preparation with <400 megabases generated. The majority of reads classified as bacterial in all samples sequenced. The proportion of reads classified as human was slightly higher than with the *C. trachomatis*/*N. gonorrhoeae* NAAT-negative spiked urine, although <29% in all but one case (Figure S6A). ≥94% of the *N. gonorrhoeae* genome was covered at ≥10-fold in 3 of 4 samples spiked at ≥10^4^ CFU/ml, suggesting that although a greater proportion of the sequencing reads were classified as human it is still possible to get both good breadth and depth of coverage of the *N. gonorrhoeae* genome in the presence of co-infection and inflammatory cells within the urine (Figure S6B). Bases classified as *C. trachomatis* were observed 12/18 of the Chlamydia NAAT-positive spiked urine DNA extracts (derived from all 3 original urine samples, Figure S6C). However, sufficient sequence data to cover the majority of a *C. trachomatis* reference genome was only obtained in 5/18 extracts all from the same original urine sample (Figure S6D). Additional species, including *Acinetobacter spp*. and *Streptococcus pseudopneumoniae*, were identified in all three urines at high proportions (Figure S7), which likely hindered recovery of *C. trachomatis*.

### Sequencing speed

Figure 2 shows the sequencing time taken to achieve an estimated given fold coverage of the reference genome for the 18 *N. gonorrhoeae* and *C. trachomatis* NAAT-negative urine samples spiked with *N. gonorrhoeae* that were depicted in Figure 1. For samples spiked with ≥10^4^ CFU/ml >20-fold coverage was typically achieved in ≤4 hours from starting sequencing.

**Figure 2.**
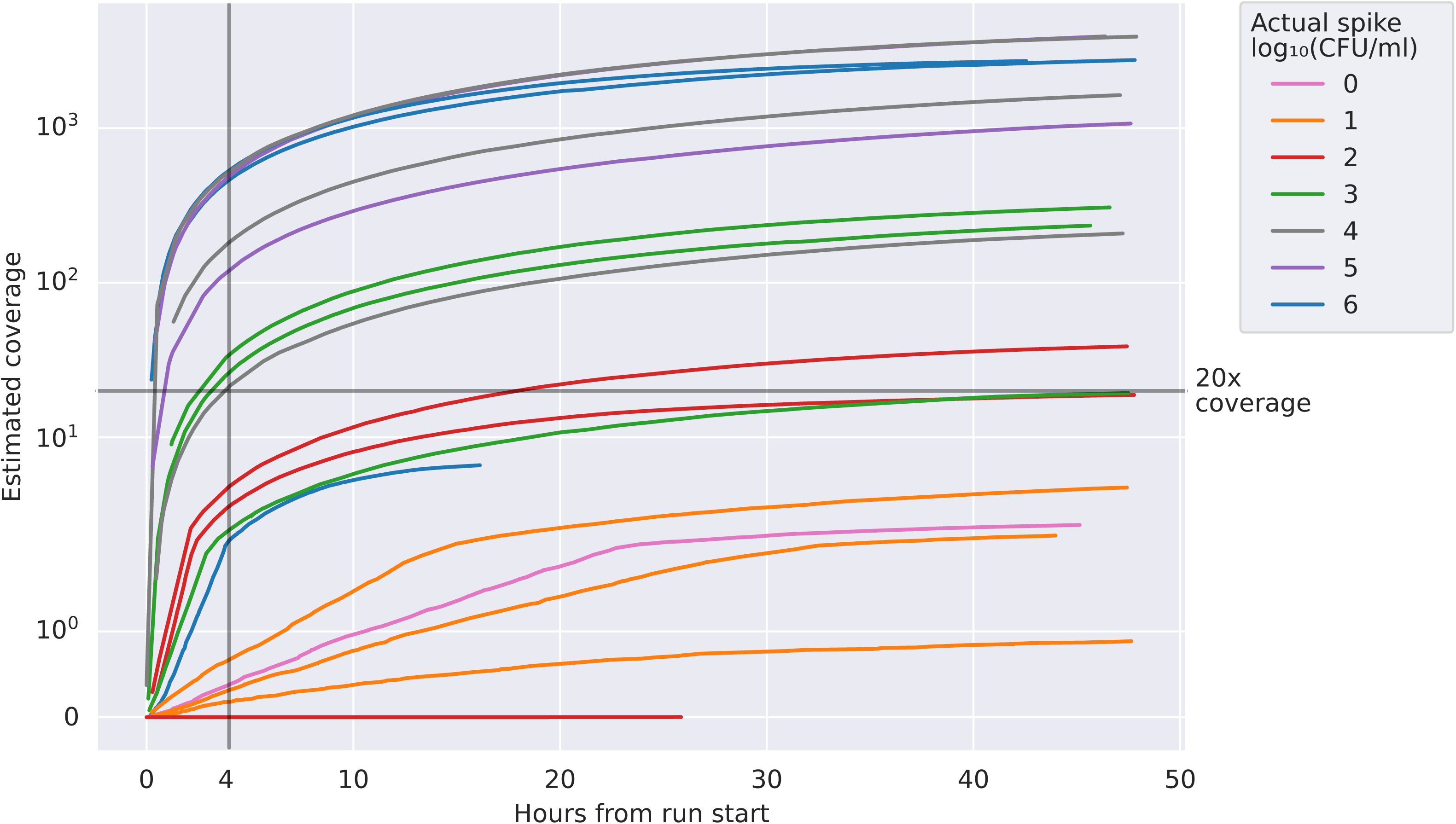
Sequencing speed in 18 *N. gonorrhoeae* and *C. trachomatis* NAAT-negative urine samples spiked with *N. gonorrhoeae*. The actual spiking concentration achieved is rounded to the nearest order of magnitude for the purposes of the legend. Estimated coverage is calculated as the number of bases of *N. gonorrhoeae* DNA sequenced divided by the length of the reference genome.

### Performance in *N. gonorrhoeae* infection

We extracted and sequenced DNA from ten individual *N. gonorrhoeae* NAAT-positive urine samples using the QIAamp kit with and without prior human DNA depletion with saponin. All samples sequenced contained detectable *N. gonorrhoeae* DNA. In contrast to simulated infection, samples processed without saponin had higher yields of bacterial DNA (Figure 3) and *N. gonorrhoeae* DNA (Figure 4). Without saponin, the number of bases per sample classified as *N. gonorrhoeae* ranged from 2.6×10^7^ to 8.8×10^8^, representing between 4% and 81% of total bacterial bases and corresponding to a median (IQR) [range] *N. gonorrhoeae* genome coverage breadth of 99.0% (98.5-99.2%) [92.8-99.4%] at a by-sample-mean depth of 76 (26-192) [11-384] (Figure 4). With saponin, the number of bases classified as *N. gonorrhoeae* ranged from 7.8×10^3^ to 1.8×10^8^, representing <0.01% to 41% of total bacterial bases. The median (IQR) [range] *N. gonorrhoeae* genome coverage breadth was 5.9% (2.0-22.2%) [0.3-98.9%] at an average depth of 3 (1-5) [1-77]. Therefore, while saponin was effective a reducing the percentage of bases classified as human from a median 67% to 46% (Figure 3) this was not sufficient to offset the detrimental effect on *N. gonorrhoeae* in clinical samples.

**Figure 3.**
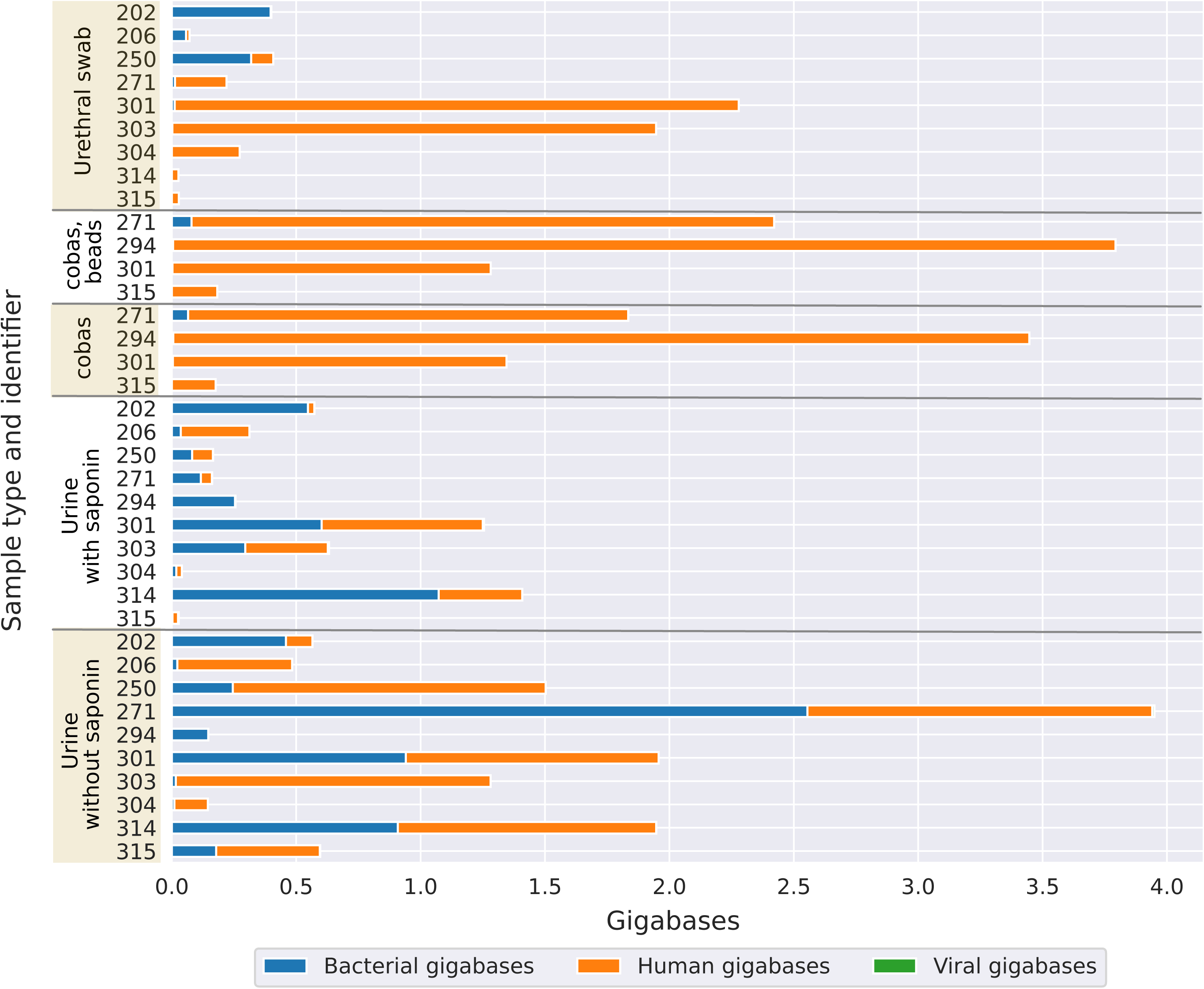
Performance in clinical samples positive for *N. gonorrhoeae*: yield of bacterial, human and viral DNA sequenced. Results from 10 participants are shown, including where available reads obtained from a urethral swab, a urine sample collected into a cobas lysis buffer (processed with and without mechanical lysis with beads), and a urine sample collected in a universal container with a boric acid additive processed with and without treatment with saponin. The plot shown was generated using sequence data demultiplexed with Porechop, after which human reads were securely deleted. To reduce the proportion of reads without an assigned barcode arising with Porechop, samples were re-basecalled and demultiplexed using Guppy for subsequent analysis.

**Figure 4.**
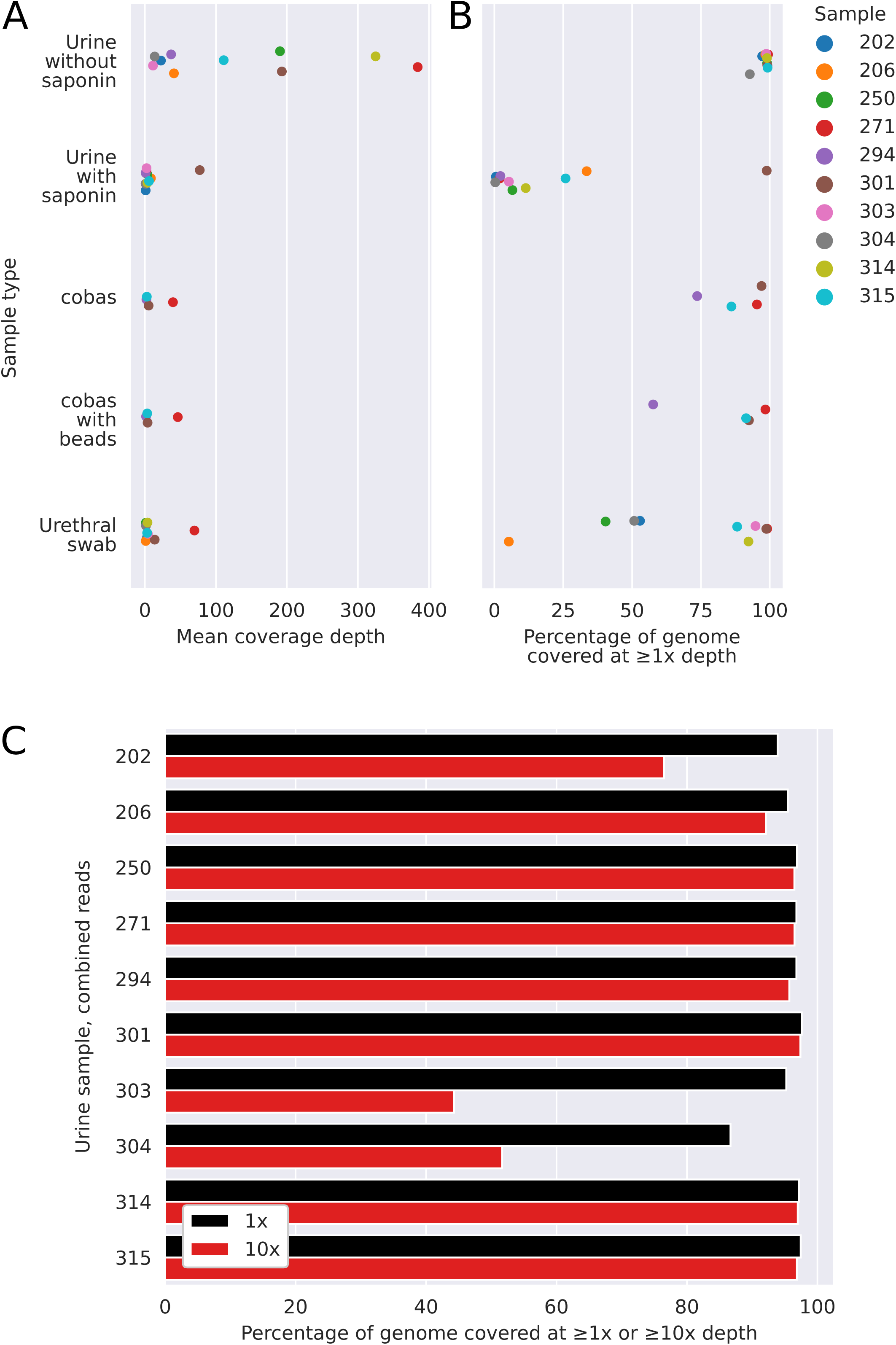
Performance in clinical samples positive for *N. gonorrhoeae*: coverage breadth and depth. Panel A shows the mean coverage depth achieved for samples processed by one the five methods tested, i.e. the total number of bases of sequence generated divided by the length of the NCCP11945 *N. gonorrhoeae* reference genome. Panel B shows the proportion of the NCCP11945 *N. gonorrhoeae* reference genome covered by at least one read. Panel C uses combined data from urine samples treated with and without saponin to represent the NCCP11945 *N. gonorrhoeae* reference genome coverage from sequencing a single urine sample on a flowcell.

Figure 5 shows the most commonly identified species across the five sample types (urethral swabs, cobas sample tubes with and without mechanical lysis with beads, and urine samples with and without saponin treatment). For 9 of the 10 urine samples processed without saponin treatment *N. gonorrhoeae* was the most abundant species sequenced, with reads classified as *N. lactamica* likely representing taxonomic misclassification of reads from *N. gonorrhoeae*. Samples processed with saponin showed differential depletion of *N. gonorrhoeae* relative to other bacteria.

**Figure 5.**
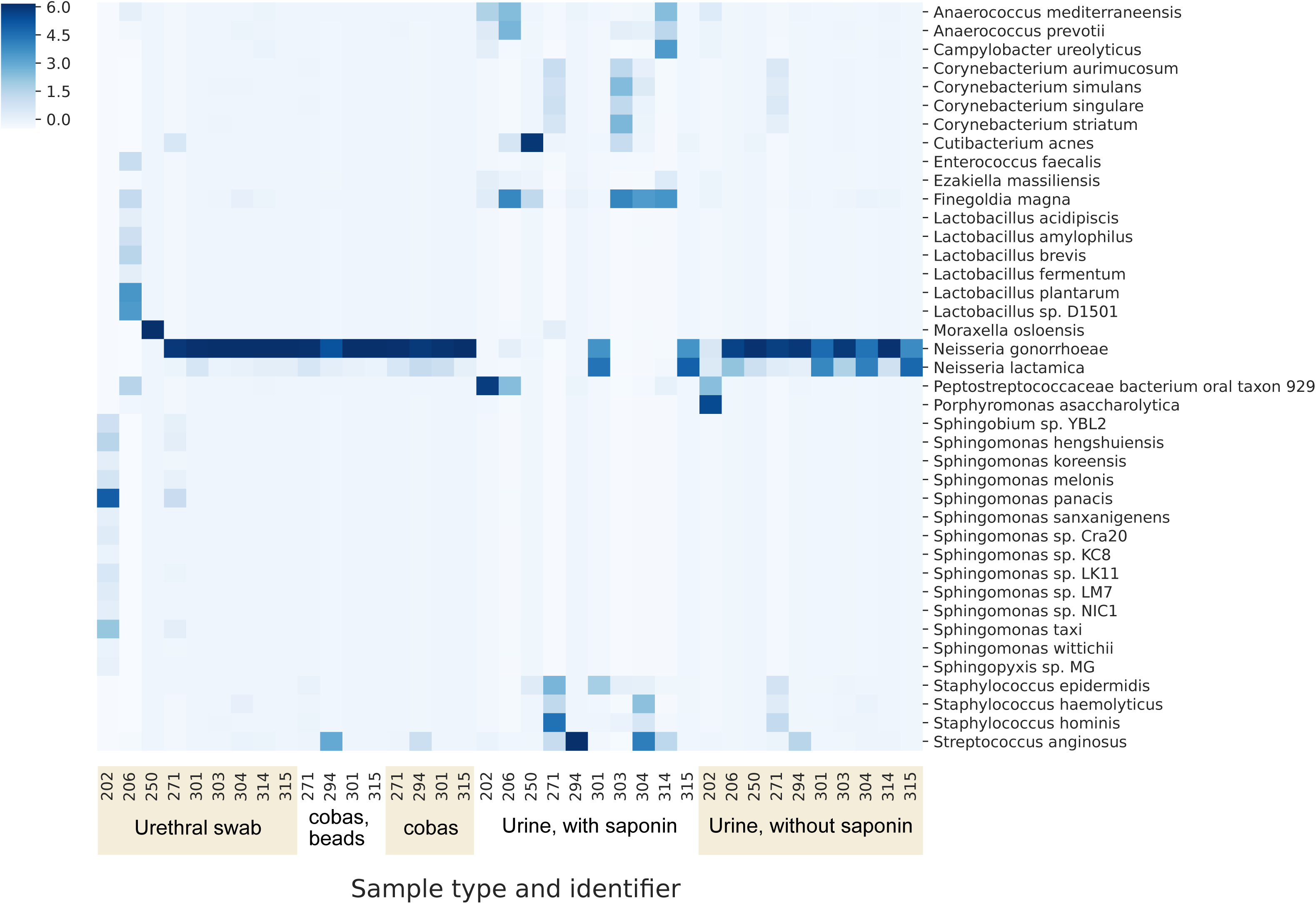
Performance in clinical samples positive for *N. gonorrhoeae*: relative proportions of species sequenced per sample. The z-score, denoted by shade, for each taxon is the number of standard deviations above the mean number of bases per taxon for each sample.

The large majority of bacterial reads were from *N. gonorrhoeae* in most urethral swab and cobas tube samples (Figure 5). However, these samples contained lower amounts of total bacterial DNA sequence (Figure 3), resulting in fewer sequenced *N. gonorrhoeae* bases and only limited coverage of the reference genome (Figure 4). Sequencing direct from cobas PCR Media tubes, without mechanical lysis with beads, the number of *N. gonorrhoeae* bases ranged from 3.7×10^6^ to 8.7×10^7^, representing between 58% and 94% of total bacterial reads and resulting in genome coverage from 74% to 97% at a per-sample-mean depth from 2 to 40. Results with mechanical lysis were similar. Similarly, the number of bases classified as *N. gonorrhoeae* from urethral swabs ranged from 1.7×10^5^ to 1.6×10^8^ representing between <0.01% and 91% of the total bacterial bases. The median (IQR) [range] genome coverage breadth was 88.2% (50.1-94.8%) [5.3-99.1] and depth 3 (1-4) [1-70].

To provide a conservative approximation of the sequence data generated by running a urine sample on a single flow cell we pooled reads obtained from the same flow cell from sequencing the same urine sample processed with and without saponin (Figure 4C). At least 87% of the reference genome was covered by at least 1 read in all 10 samples and ≥95% of the reference genome in 8 samples. In 7 samples ≥92% of the reference genome was covered at a depth of ≥10-fold. We explored predictors of successful sequencing (Table S2). Within the limits of the small sample, there was no relationship between the percentage of the reference genome with ≥10-fold coverage and NAAT CT values, the time between collection and processing, the concentration of DNA loaded on to the flowcell post amplification, or the number of active flowcell pores at the start of sequencing (all Spearman’s rank p>0.2). In the 3 samples that did not achieve ≥92% coverage at 10-fold depth, one had high levels of contaminating bacteria DNA, sample 202 (*Porphyromonas asaccharolytica*, Figure 5). In another, sample 303, the proportion of human reads was higher than in other samples, e.g. without saponin, 99%, compared to a median (IQR) of 67% (51-90%). The final sample, 304, had a relatively poor sequence yield overall, the barcode without saponin yielding only 0.1 Gb of data, compared to a per-barcode median of 1.4 (0.7-2.0) Gb.

## Discussion

We demonstrate it is possible to extract sufficient quantities of *N. gonorrhoeae* DNA directly from urine samples from men with symptomatic urethral gonorrhoea and to sequence to achieve near complete reconstruction of the *N. gonorrhoeae* genome. It was possible to achieve coverage of ≥87% of the genome in all ten patient samples, and ≥92% coverage breath at ≥10-fold coverage in 7 (70%) samples. Through simulated infections we demonstrate that if *N. gonorrhoeae* is present at ≥10^4^ CFU/ml sequencing of the large majority of its genome can be frequently achieved.

Our initial experiments with spiked *N. gonorrhoeae* NAAT-negative samples showed the four DNA extraction methods tested were broadly comparable and potentially any could be applied for sequencing direct from urine samples. However, we obtained contrasting results from attempted human DNA depletion in simulated and actual infections. In simulated infections, freshly cultured *N. gonorrhoeae* was spiked into urine samples and DNA extracted; in this setting both saponin and the MolYsis kits improved *N. gonorrhoeae* DNA yields by differentially depleting human DNA, the former to a greater extent. However, in actual infections, *N. gonorrhoeae* appears to have been more susceptible to lysis by saponin, possibly reflecting damage to *N. gonorrhoeae* cells in storage, transport and by host inflammatory cells. This resulted in higher yields of *N. gonorrhoeae* DNA when saponin treatment was omitted.

The depth and breadth of coverage of the *N. gonorrhoeae* genome achieved in the majority of samples spiked with ≥10^4^ CFU/ml of *N. gonorrrhoeae* and the *N. gonorrrhoeae*-positive clinical samples should make detection of antimicrobial resistance determinants possible, as well as comparisons of genomes for transmission tracking. However, the relatively high per base error rate of Nanopore sequencing means specific bioinformatic approaches are required to produce a consensus genome without an unacceptable number of false variants. This follow up work is an area of active research at present.

Initially when planning the study, we considered that contamination with human DNA would be the principal technical challenge to be overcome. Although the clinical samples from *N. gonorrhoeae* infection tested still contained human DNA, in the majority of samples sufficient *N. gonorrhoeae* DNA was present for successful sequencing without specific human DNA depletion. However, large amounts of human DNA present in one clinical sample resulted in reduced genome coverage. Presence of high levels of other bacterial species impaired the yield of pathogen DNA in simulated infections and in one of the samples from clinical *N. gonorrhoeae* infection. For simulated infections this is likely because this study relied on samples discarded after routine testing. As such, these samples had typically spent several days at ambient temperature in transport and in the laboratory, which allowed time for bacterial overgrowth to occur. This same issue was present in study participant samples, albeit to a lesser extent, as samples from Brighton were couriered overnight at ambient temperature to Oxford. The large amounts of DNA from other bacteria in one of the clinical samples occurred despite the use of boric acid as an additive to reduce growth. We also tested if collection of urine directly into an unselective cell lysis buffer would prevent bacterial over-growth. This approach was successful in preventing contamination with other bacterial DNA, however it also better preserved human DNA, such that the total amount of bacterial DNA sequenced, and hence yield of *N. gonorrhoeae* DNA was lower using this approach. *N. gonorrhoeae* DNA yields from urethral swabs were also low, which may represent low numbers of organisms collected, particularly as obtaining these swabs required a second urethral swab (in addition to that taken for routine culture), which was potentially uncomfortable for participants.

Although our results provide a proof of principle, the applicability of sequencing in its current form is also limited by the time taken to prepare samples for sequencing, this requires up to 10 hours, largely due to the need for prolonged PCR amplification of very low quantities of input DNA. Current reagent costs are also >$400 per sample; while these could be reduced by multiplexing multiple samples per flow cell this would reduce sensitivity.

Our results also highlight another current limitation of metagenomic sequencing, the potential for contamination, particularly as the approach relies on non-selective amplification of all DNA present. In one of our seven negative control urine samples around 15% of the *N. gonorrhoeae* reference genome was covered at high depth, however the coverage was very uneven (Figure S5). This partial coverage of the reference genome may have arisen from contamination with PCR amplicons. This reinforces the need for metagenomic sequencing based studies to include appropriate negative controls. Additionally, confirmation of the presence of *N. gonorrhoeae* may also require achieving coverage of a substantial proportion of the reference genome, and further metagenomic sequencing studies of patients with and without *N. gonorrhoeae* infection are required to assess this and determine thresholds for robustly identifying infection.

The focus of this manuscript was to optimize laboratory methods, which we have successfully achieved. This work provides a firm foundation for developing bioinformatic methods for confirming the presence of *N. gonorrhoeae* and resistance gene identification using Nanopore data. If this can be achieved, same-day metagenomic diagnosis of gonorrhoea infection and antimicrobial resistance is likely to be possible.

## Acknowledgements

The authors thank the microbiology laboratory staff of Oxford University Hospitals NHS Foundation Trust and the Royal Sussex County Hospital, Brighton for providing assistance with sample collection.

## Declaration of interests

JO has received consumables, research funding and conference expenses from Oxford Nanopore Technologies. No author has a conflict of interest to declare.

## Funding

This work was funded by the Centers for Disease Control and Prevention, under Broad Agency Award FY2018-OADS-01.

